# Listing candidate diagnostic markers and transcriptomic exploration of the molecular basis of a type of male infertility (Non-Obstructive Azoospermia), via next generation sequencing methods

**DOI:** 10.1101/778670

**Authors:** Balagannavar Govindkumar, Basavaraju Kavyashree, Akhilesh Kumar Bajpai, Sravanthi Davuluri, Kannan Shruthi, SS Vasan, M Madhusudhan, S Chandrasekhar Darshan, Chitturi Neelima, Balagannavar Vashishtkumar, Oguru Sailaja, K Acharya Kshitish

## Abstract

Studying the molecular basis of Non-Obstructive Azoospermia (NOA), a type of male infertility with failed spermatogenesis at various stages, can also help in exploring molecular basis of human spermatogenesis and possibly pave way to identify new targets for male contraceptive development. Hence, we initiated a functional genomics study by applying RNA-seq. Testicular biopsies collected from donors with Non-Obstructive Azoospermia (NOA), Obstructive Azoospermia (OA), Congenital Bilateral Absence of the Vas Deferens (CBAVD), and Varicocele (VA) conditions. Strong association of 100+ genes with human spermatogenesis and NOA has been detected via NGS-based transcriptomic analysis. In addition, 20 RNA molecules have been short-listed for potential diagnostic applications (non-obstructive azoospermia vs. obstructive azoospermia, varicocele or normal). A hierarchical list of several genes and alternatively spliced mRNAs, transcribed differentially in NOA, is reported - based on a ‘strength of association’. Such association with NOA, spermatogenesis or both is a new finding for many genes as revealed by a comparison with a newly prepared comprehensive list of genes having such association with human spermatogenesis/NOA. Many top-ranking genes involved in viral gene expression were up-regulated in testes from NOA-patients, while those associated with an antiviral mechanism were down-regulated. A tangential finding: while most well-established control mRNAs did not qualify, two new ones worked best in RT-qPCR experiments. Needle-aspiration of testicular biopsies, followed by the use of short-listed promising candidate biomarkers (i.e., 16 mRNA & 4 chimeric transcripts) and control mRNAs in RT-qPCR-based diagnostic assays, may help to avoid open surgeries in future.

## Introduction

Male infertility seems to be increasing in recent years (1, 2) and azoospermia is one of the common causes with two main subtypes: Obstructive Azoospermia (OA), where there seems to be a ductal obstruction while spermatogenesis is normal, and NOA, where the spermatogenesis is impaired. NOA is very difficult to deal with clinically, and testicular surgical biopsy is the only way to detect the disorder (3). Detailed studies are needed to understand the genetic and molecular basis of the NOA condition, as well as short-listing the genes, or even the corresponding mRNA isoforms, that may have a strong association with this disorder.

Compared to the need to address NOA, controlling the growth of the human population is a contrasting challenge faced by multiple countries - particularly India and China. The family planning programs can be helped by the development of reversible and easy-to-use male contraceptives. There have been efforts to find new contraceptives for use by men. But there is a need for further efforts in this direction. Understanding the molecular basis of spermatogenesis is a common pre-requisite for male-contraceptive-development as well as addressing NOA. Most men suffering from NOA seem to have completely normal general physiology and sexual desire (4). Hence, understanding the molecular mechanism of spermatogenesis by using NOA as a case study can provide an excellent opportunity to (a) not only discover potential molecules for further research in diagnostics, prognostics, and therapeutics, but also (b) identify candidate target molecules for research towards the development of new male contraceptives.

It is not surprising that globally, researchers have tried to enhance the understanding of the molecular basis of spermatogenesis. It is clear that specific patterns of expression of the genes are essential for spermatogenesis, and several studies have been conducted particularly at the transcriptional level (e.g., 5, 6, 7, 8). Even though there have been improvements in studying transcriptomics (9), not all genes essential for spermatogenesis and those associated with NOA have been identified, and more studies are required on various fronts of gene expression associated with spermatogenesis and infertility in men (e.g., 10). Realizing the potential use of screening expression of most of the genes, and identifying genes associated with NOA via differential expression profiling, many scientists used the microarray techniques (e.g. 11, 12, 13). Such microarray-based studies on human testes and many other studies carried out on the mouse model have certainly enhanced the knowledge base of the transcriptome of the testis in normal and NOA conditions. Such studies have also helped us understand the molecular basis of the spermatogenesis as well. However, while microarrays form a phenomenal technique that also contributed significantly to the transcriptomics of several conditions of interest across species, such gene expression profiling approaches have several limitations (14). Despite the advantages offered by the Next-Generation Sequencing (NGS) methods (15), which have been applied in genomic contexts of NOA (e.g., 16, 17), RNA-sequencing has not been applied to thoroughly explore the transcriptomic differences of the testis tissue in NOA compared to normal samples. RNA-seq approach has been taken up to explore the transcriptomic of semen samples (18). Testicular microRNA sequencing has been performed using NGS methods recently, albeit for 3 NOA samples only (19). One of the key reasons for lacunae in understanding the molecular basis of human spermatogenesis is the limitation in the availability of testis biopsy samples. That is why, even in cases such as NOA, where biopsy-based examinations are permitted, there have not been enough studies with a good number of samples. Recently single-cell RNA sequencing has been used to explore the transcriptome of specific cell types of testicular cells where tissue sample from a single NOA-donor has been used (20).

There are two important reasons why mRNA-sequencing (RNA-seq) of testis samples from a reasonable number of NOA patient-donors needs to be undertaken: Many genes were not considered in the older microarray-platforms utilized by earlier studies, and RNA-seq experiments provide an opportunity to address ‘all’ genes in the human genome. Also, RNA-seq has the potential to identify alternatively spliced transcript isoforms that may be associated with the disorders of interest.

Hence, we took up efforts to establish NGS-based transcriptome of the testis samples from NOA-patients and compare it with control conditions. Apart from the transcriptomic analysis, the study also addresses another major gap in the efforts to understand the molecular basis of the disorder, viz., the lack of compilation of genes already associated with NOA and/or human-spermatogenesis in general, and listing all such genes based on their strength of association.

## Material and methods

### Sample collection and RNA isolation

Testicular biopsies were collected from 18, 7, 2 and 2 donors, respectively, with Non-Obstructive Azoospermia (NOA), Obstructive Azoospermia (OA), Congenital Bilateral Absence of the Vas Deferens (CBAVD), and Varicocele (VA) conditions, following ethical procedures approved by the Institutional Biosafety Committee (IBSC) at Institute of Bioinformatics & Applied Biotechnology (IBAB) and Ankur-Manipal hospital, Bengaluru. The age of donors ranged from 28 to 38 years with a mean of 32.2 and a standard deviation of 4.15. The samples were stored in RNA-later solution (Ambion, cat # AM7024), according to the manufacturer’s guidelines. RNA was extracted using the RiboPure kit (AM1924) and quality was checked with formaldehyde agarose gel electrophoresis as well as a micro-volume spectrophotometer.

For NGS, testicular RNA from 8 NOA, 2 OA and 2 Varicocele donors were used. In addition, 2 normal commercial RNA (cat. No.: 636533, Asian: lot No: 1105041; Caucasian lot. No.: LOT1105214A) samples were used.

### Sequencing and analysis

Paired-end library preparation was carried out using TruSeq RNA Library Prep Kit v2, and sequencing was done using Illumina HiSeq 2000. The sequencing depth per sample ranged from 43 to 73 million reads and the read length was 100bp. With an intention of increasing the number of controls, RNA-seq data sets for normal human testis with reasonable depth (>40M) were identified in SRA and downloaded (see Supplementary Table M1). The quality of selected and downloaded raw data corresponding to nine samples was found to have a mean of Phred quality score ranging from about 27 to about 37. FastQC (https://www.bioinformatics.babraham.ac.uk/projects/fastqc/) tool was used to assess the quality of reads. Reads with a minimum mean quality score of 20 or lower, and or adapter content of 9 or higher percentage were trimmed using seqtk (https://github.com/lh3/seqtk) tool. Alignments, identification of transcripts and the chimeric/transplice molecules, and their quantification were performed by Kallisto software (21). Human transcript (GRCh38.90) sequences from Ensembl were used as a reference. Differential expression analysis was done using the Bioconductor package limma (22). To test the heterogeneity among samples, they were hierarchically clustered based on the expression values of transcripts across samples, using an R package (ggplots, 23). The clusters were visualized using RColorBrewer (http://colorbrewer.org).

Apart from the standard data analysis procedures for identifying transcripts expressed differently in NOA, a novel scoring system (http://www.startbioinfo.com/methods/StA) has been applied to rank the transcripts based on the ‘Strength of Association’ (StA) with NOA. The StA score depended on the P-value, fold change and the number of contradictions across groups. Up- and down-regulated transcripts were hierarchically arranged based on their StA scores.

### RT-qPCRs

We tried to select about 80 transcripts among the top-scoring transcripts from the StA-score based hierarchical list, for further validation using RT-qPCRs on more samples. Forty transcripts, where unique regions were available for primer-designing, were selected among up-regulated transcripts and, similarly, 40 transcripts were also selected among the down-regulated transcript-list. Primers were designed for each of these 80 transcripts. The effort was made to pick at least one of the primers for each transcript from an exon-exon junction. An in-house unpublished version of the ExPrimer tool (24) was used to design primers. RNA extracted from 17 NOA, 7 OA, 2 Varicocele, 2 CBAVD, and 2 commercial normal samples were used. One of the NOA samples used for RNA-seq could not be used in RT-qPCR due to insufficient quantity. cDNA was prepared using ‘ImProm-II™ Reverse Transcription System’ (Promega cat No: A3800) according to the kit protocol. RT-qPCR experiments were performed using Sybr green (Kappa cat no: KK4601) reaction mixture, using the manufacturer’s protocol, and the StepOnePlus instrument of Applied Biosystems, or the Rotor-Gene Q of Qiagen.

### Selection of control genes for RT-qPCRs

Control genes were carefully selected. RNA-seq data were screened to test the suitability of 10 commonly used control genes, such as GAPDH and actin, and their corresponding transcripts. Most of the transcripts of these genes were found to be differently transcribed in NOA compared to normal. Hence, the following criteria were used for the selection of better control transcripts: a) the P-value should be high for differential expression (≥0.4); b) variation of TPM values should be minimum across the groups. A total of 3 control genes, each with multiple product-sizes corresponding to each transcript tested, were used. All three transcripts corresponding to the selected control transcripts were first confirmed to be undifferentiated by preliminary RT-qPCR experiments. The product sizes were kept between 90 to 280 bp. But the size was kept similar for products within an experiment. The primers were designed keeping this in mind while ensuring coverage of unique-regions in the test transcripts.

RT-PCR was first carried out for all 80 transcripts with 1 control and 3 NOA samples. No amplifications were observed in any of the samples, despite attempts with varying annealing temperatures, for 12 transcripts. RT-qPCRs were carried out for the remaining transcripts in two rounds: 1) These transcripts were tested first with 7 NOA, and 1 each of normal, OA and varicocele samples. Only the transcripts with expression patterns similar to those observed via RNA-seq experiments were chosen for the next round. 2) In the second round, we performed RT-qPCR using 1 normal, 2 CBAVD, 6 OA, 1 varicocele and 10 NOA samples.

The 2^-ΔΔC^T method (25) was used to identify the fold change in the expression level of transcripts in NOA compared to the control conditions, and t-test was used to assess the significance of the difference. For t-test, first, the difference was noted between the Ct-values of each test transcript in a sample and the C_t_ value of each of the control transcript per sample. Since two control transcripts were used for each test transcript, the mean of two such differences in C_t_-values (mean-delta-C_t_) across control transcripts was calculated per sample. Finally, we estimated the significance of the difference in the mean of such mean-delta-C_t_ in all NOA samples vs. all control samples. The RT-qPCR results were represented as box plots and it was generated using the Bioconductor package (ggplot2).

### Functional analysis

Enrichment of functions among the top 3,000 each of up- and down-regulated genes was performed by identifying Gene Ontology (GO) (26, 27) Terms and pathways over-represented among each set using DAVID (https://david.ncifcrf.gov/gene2gene.jsp). This was repeated with the top 250 each of the same two sets of genes.

To obtain an indication of the relative extent of studies of each of the top 250 protein-coding genes identified to be up- and down-regulated in NOA in the current study (top 500 genes), the existing information about these genes was obtained using the following methods: a) Functional summary information was collected from NCBI-gene, Gene Cards and UniProt (via DAVID, https://david.ncifcrf.gov). b) Obtaining the number of articles where the gene name appears in likely association with spermatogenesis or NOA: Literature searches were performed using Annokey (28). Two types of searches were performed where the gene name was used along with one of the following terms: ‘non-obstructive azoosperm*’ and ‘sperm*’. c) In addition, to find out the general extent of studies carried out on each gene, GenCLIP (29) was used. To be sure of the known status of top-genes, a parallel effort to biocurate was also made to compile a set of genes associated with spermatogenesis and/or NOA by carefully designing a set of terms and phrases, for searching in PubMed.

To find GO terms associated with spermatogenesis for each gene, the terms were first downloaded from Biomart (http://asia.ensembl.org/biomart/martview/), specific terms of relevance were carefully selected and Java scripts were written to screen up to about 15,000 terms and detect the pre-identified relevant terms in each of the 500 rows (corresponding top 250 up- and down-regulated genes).

## Results

Quality of the RNA extracted from testis samples and the quality of reads obtained from RNA-seq were found to be good. The sequencing depth was also good across samples (a mean of 56 million reads).

### Clustering analysis

The analysis showed that the overall gene expression profiles in Non-Obstructive Azoospermia (NOA) were different compared to control samples with normal spermatogenesis - irrespective of sub-types within each set. More importantly, the observations supported the decision to group normal testis-samples with those from other conditions where spermatogenesis occur normally, viz., Varicocele (VA), Obstructive Azoospermia (OA) and Congenital Bilateral Absence of the Vas Deferens (CBAVD). The grouping not only enhanced the number of control samples, but also helped to eventually identify candidate markers that are more unique to NOA.

### RNA-sequencing

The raw RNA-seq data are available at Sequence Read Archive (SRA) database. When the mean expression level of alternatively spliced transcripts across NOA samples was compared with the corresponding levels in control samples, 2,201 transcripts were found to be up-regulated in NOA, and 16,363 down-regulated, with P-value and log2-fold change of <0.05 and >=1.5, respectively. Thus, there were substantially more down-regulated transcripts compared to up-regulated ones. This is probably because of the absence of, and/or a very low number of, germ cells in NOA-testis, particularly the post-meiotic cell types.

It is a common practice to short-list genes via functional analysis after applying thresholds on the fold-change and statistical significance of the difference in the expression levels. This popular approach of finding genes associated with a disease may generate a bias towards well-studied genes, and leads to underestimation of the consistency of association of genes with the disease of interest. Hence, we adopted a ‘Strength of Association’ (StA) scoring method that allows a hierarchical listing of molecules based on consistency in differential transcription. As mentioned below, multiple observations suggest that the StA scores may be useful indicators of the reliability of gene expression patterns.

The differential transcription was found to be more prominent and reliable among the down-regulated transcripts than among up-regulated ones. For example, the StA score of the top 100 transcripts of the hierarchical up-regulated list was >79.1, while the same was >91, for the down-regulated list. The lower reproducibility among up-regulated transcripts was further supported by the results of the RT-qPCR experiments - where the top up-regulated mRNAs showed poor agreement across samples, unlike the top down-regulated ones.

### RT-qPCR experiments

Interesting observations were made while selecting control genes/transcripts for RT-qPCR experiments. Among the 10 commonly reported/used house-keeping genes screened based on differences in the mean TPM values obtained from the RNA-seq experiments, and the corresponding statistical reliability of the difference, only one gene, HPRT1 appeared to be a reasonably reliable control gene for RT-qPCR experiments. It had a difference of 32.9 in mean TPM values across conditions (103 TPM in normal vs. 136 in NOA), coefficient of variation (CV) of 0.54 and a P-value of 0.40. Hence, we tried choosing controls from the list of undifferentiated transcripts obtained from the current RNA-seq data. The difference in the level of expression between NOA and control conditions, in terms of mean TPM values, was 0.14 for one chosen transcript and 0.89 for the other, and the corresponding P-values obtained after t-tests, were 0.91 and 0.9, respectively. The mean TPM value across all samples and groups, and the corresponding CV value were 24.19 and 0.16, respectively, for the first transcript, and 103.15 and for the other. Preliminary RT-qPCR experiments confirmed that all 3 control genes have minimum variation across samples. Thus, we discovered two most reliable control transcripts for use in studies comparing gene expression in human testis across NOA, OA, varicocele and normal conditions.

Results were compiled across two rounds of RT-qPCR experiments with a total of 17 NOA, 2 normal, 2 Congenital Bilateral Absence of the Vas Deferens (CBAVD) and 7 Obstructive Azoospermia (OA) & 2 varicocele (VA) samples. Each round included at least one each NOA and control RNA samples that were used for RNA-sequencing. None of the down-regulated transcripts completely contradicted the NGS-based expected trend of differential transcription, though some of the individual samples showed contradictory trends for some transcripts. More contradictions were seen among up-regulated transcripts. Overall, a high concurrence between the results from NGS vs. RT-qPCR should not be expected perhaps given the potential variations in the reads mapping to different regions of transcripts, particularly to the unique regions from where the primers were designed. Hence, it is not surprising that only 16 of the down-regulated transcripts, and none of the up-regulated ones, showed 100% consistency in differential expression. Up-regulated transcripts had lower StA score to begin with.

The P-value for the significance of difference in the delta Ct values of the 16 transcripts that were consistently down-regulated in NOA ranged from 1.00E-9 to 1.40E-05, while the (log2) fold change of these transcripts ranged from 3.55 to 7.63 (Fig. 1). These transcripts seem to be more promising biomarker candidates for differentiating testes samples with NOA from other conditions where spermatogenesis is occurring normally - irrespective of the possible sub-types in NOA or the control conditions. When the individual data points (mean delta-CT values) for the normal condition are compared with NOA, many seem to overlap in Figure 1. But this was not a concern because no such overlaps were seen in individual experiments in any case. This is illustrated using data from individual experiments for 3 selected transcripts that showed the least significant P-values across all experiments. The observations support the StA scoring method where one of the criteria was that the highest level of expression for a down-regulated transcript should be lower than the lowest value of that transcript in normal condition. RT-qPCR results did not show consistent agreement across all tested samples for the up-regulated transcripts.

**Figure 1.**
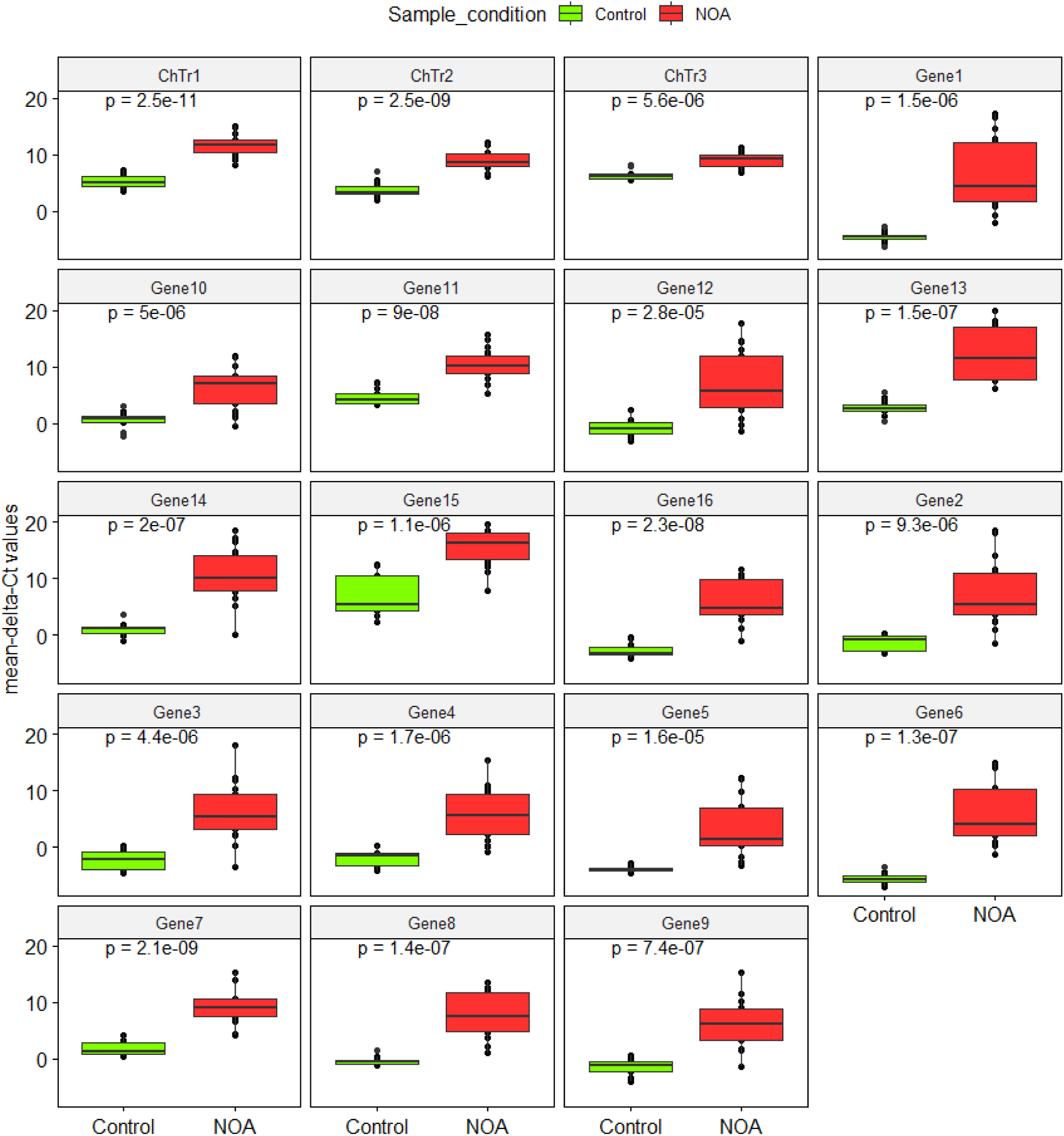
Results of RT-qPCR experiments indicating the relative amount of selected transcripts across control and NOA samples. The delta Ct values represent the mean difference in the Ct-value of each transcript with each of the control transcript, across NOA vs. control samples. The numbers within each box plot indicate P-values obtained following the t-test.

NGS data analysis showed 8 chimeric transcripts (ChTr) of which six were down-regulated and two up-regulated in NOA. Three of the down-regulated chimeric transcripts showed consistent down-regulation in NOA during RT-qPCR-based validations (Figure 1). Hence, these 3 chimeric transcripts were also considered to be equally potent candidate markers with diagnostic applications.

### Data mining, biocuration and comparing with RNA-seq results

Using carefully designed query-sets, 313 relevant hits were obtained in PubMed from which we compiled 452 genes reported being associated with NOA and 168 genes associated with spermatogenesis. A total of 1,574 transcripts were detected among 394 differentially transcribed genes of the 452 biocurated genes. Of these transcripts, 1,272 were down-regulated transcripts and 302 were up-regulated. The analysis of the number of PubMed hits per gene in general or in the context of NOA or spermatogenesis enhanced our confidence in the list of down-regulated genes derived by RNA-sequencing and StA. Interestingly, most of the top-scoring genes were not earlier established to be associated with NOA. After combining the manual biocuration, Gene Ontology (GO) analysis, and AnnoKey-based searches, 209, 108 & 101 of the top 250 down-regulated genes were found to be newly associated with NOA, spermatogenesis & both, respectively. Similar associations were found with 238, 160, & 164 of the top 250 up-regulated genes.

The current results not only provide a relative rank for the strength of association for many genes that were already established to be associated with NOA but also specify the alternatively spliced form of these genes. Some of the important genes down-regulated in NOA that seem to be well studied in the context of NOA as well as spermatogenesis, with more than 200 research articles in general, and more than 10 articles in the context of spermatogenesis or NOA. In fact, one of these genes has another alternatively spliced transcript with a completely opposing expression pattern (up-regulated) in NOA compared to normal. There were also many well-studied genes corresponding to the top 250 down-regulated transcripts which do not seem to be recognized earlier to have an association with spermatogenesis or NOA.

Similarly, among the well-studied up-regulated genes, several genes had no hits for spermatogenesis or NOA. However, the genes up-regulated in NOA may not necessarily have an association with spermatogenesis.

### Functional analysis based on Gene Ontology (GO) terms was carried out for Biological Process (BP), Molecular Function (MF) and Cellular Component (CC)

The BP terms enriched among top 3,000 NOA-down-regulated genes include obvious biological processes related to testis functions. About 232 genes are associated with sperm development or related processes. Two such prominent processes were ‘spermatogenesis’ (count: 163 genes, Benjamini P-value or Bpv: 2.66E-42) and ‘spermatid development’ (42 genes, BPv: 1.08E-13). Broad GO terms such as multicellular organism development (112 genes, Bpv: 0.00006) that may not have significance to the list were ignored. Other specific processes that seem to be negatively affected are cilium assembly, cilium morphogenesis, and related processes, cell division and related processes such as meiosis and DNA repair, and several aspects of RNA synthesis, process and post-translational modifications (Fig. 2).

**Figure 2.**
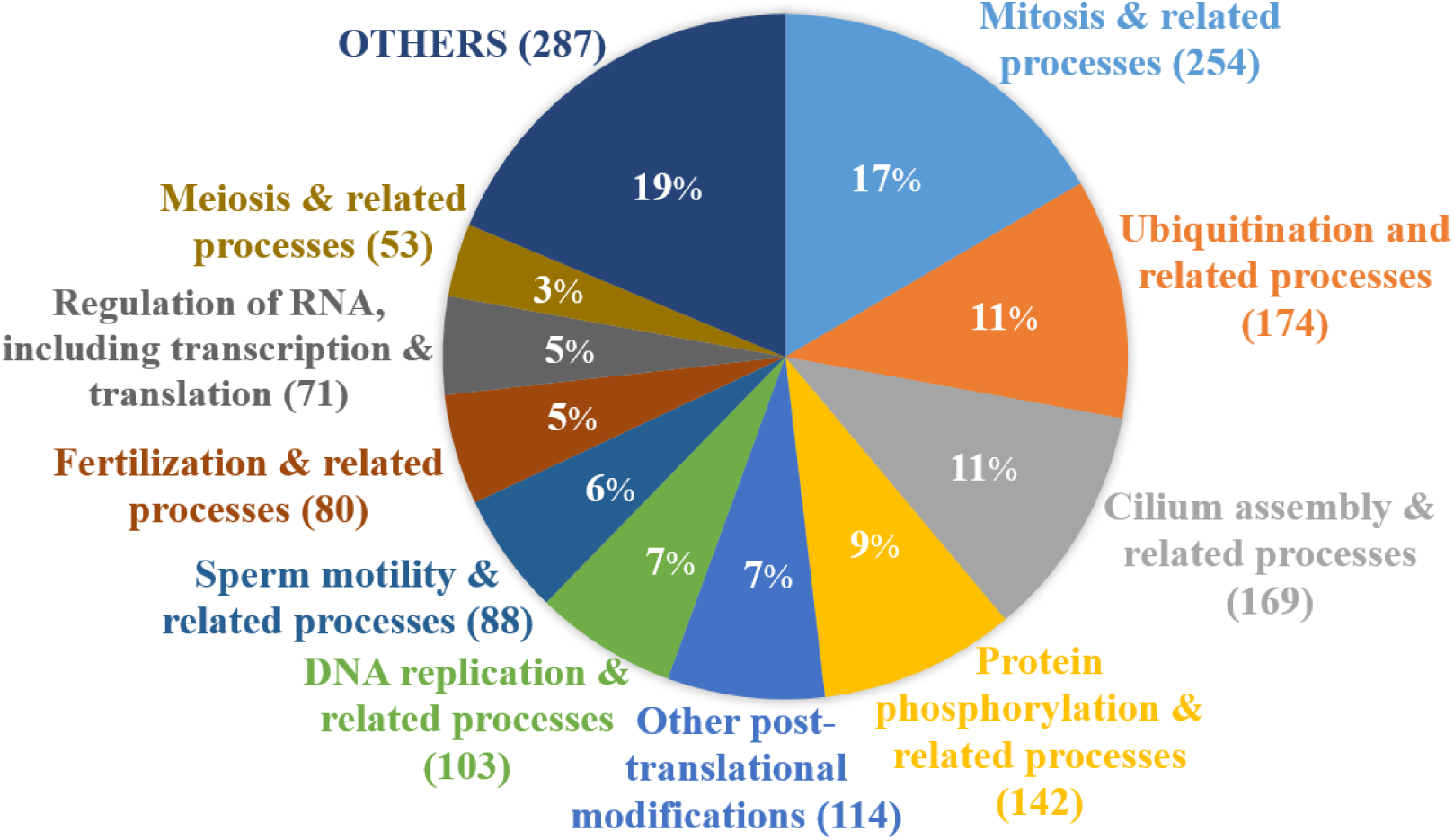
Distribution of specific functions among top 3,000 genes, after removing some of the broad processes viz., spermatogenesis and spermatid development, spermatid nucleus elongation, and cell differentiation. The numbers in the parantheses indicate the number of genes associated with the corresponding processes.

Many of the significant molecular functions, biological processes, cellular components and pathways enriched among top 250 up- or down-regulated genes were also enriched among the corresponding larger set of 3,000 genes. This observation indicated the reliability and utility of the StA-score-based ranking of differentially transcribed genes. But, there were also many GO terms and pathways that were found exclusively among the top 6,000 differentially transcribed genes as indicated below.

### Top GO terms or pathways

Interestingly, viral activity is indicated among the up-regulated genes while the antiviral mechanism is represented among the down-regulated genes. The GO terms such as ‘viral transcription’ (P-value <3.2E-22, 63 genes) and ‘viral nucleocapsid’ (P-value <0.0067, 11 genes), and the ‘viral mRNA translation’ (P-value <3.71E-31, 66 genes) pathway are enriched among the top-3,000 up-regulated genes. The top 250 up-regulated genes also showed over-representation of the ‘viral transcription’ and ‘viral mRNA translation’. On the contrary, the pathway ‘ISG15 antiviral mechanism’ (Reactome P-value <3.16E-06, 24 genes) is over-represented among top 3,000 down-regulated genes. NS1 mediated effects on host pathways, rev-mediated nuclear export of HIV RNA, nuclear import of Rev protein, and vpr-mediated nuclear import of PICs were among other significant (P-value 0.02 to 4.6E-05) pathways or GO terms enriched among down-regulated genes. These observations hint at a possibility of testicular viral infections and/or a failure to counter such infections as a potential cause of NOA. Nevertheless, more detailed studies are needed before raising such a hypothesis.

## Discussion

The current study derived the first list of genes and transcripts that are arranged hierarchically based on their <u>St</u>rength of <u>A</u>ssociation (StA) with the Non-Obstructive Azoospermia (NOA). Some of the top-scoring genes found to be associated with NOA in the current result were also reported to have such an association earlier via microarray studies are FAM71F1, CAPN11, BTG4, OAZ3, AKAP4, CHRNB3, CCDC83, PDHA2, PDCL2, ADAM29, SPATA3, SPERT, UBQLN3, SPACA4, FBXO39, GGN, H1FNT, ZCCHC13 and POU5F2. But, the study also successfully identified several new molecules, including 3 transpliced RNA molecules, associated with the NOA and human spermatogenesis, and short-listed promising biomarkers that can be explored for clinical applications.

The amount of samples collected from azoospermic patients for biopsy can be reduced if RT-qPCR based assays can replace histological examinations. There have been many efforts to find new diagnostic methods (3). RT-qPCR based assays have the potential to facilitate fine needle aspiration and avoid open surgical biopsies (30, 31, 32). Currently short-listed set of transcripts seem to be good candidates to further test for such an application. Hence, it is crucial, to further test the biomarker status of the 19 transcripts, which showed a 100% agreement across 18 NOA and 22 control samples in the current study, with more samples. The efficiency of future research in this direction, can be further assisted by utilizing the suitable control transcripts discovered in the current study.

Further research with approaches similar to the ones employed in the current study, but with a consideration of specific sub-types of NOA can help in identifying genes and transcripts that may be consistently associated with each sub-type of NOA. The discovery of novel biomarkers for each sub-type may also eventually lead to ways to distinguish NOA patients with a reasonable number of spermatids and/or sperm in the testis. Currently, the desired overall success has not been achieved in most cases with the *in vitro* germ cell injections, as evident by multiple reports of fertilization and/or pregnancies post micro-injection of round (e.g., 33) or elongated (e.g., 34) spermatids or sperm (e.g., 35) retrieved surgically from patient-testes. However, more research with culture methods combined with continued efforts to discover more precise biomarkers could help to achieve such success. Some of these biomarkers could also be candidate target molecules for research towards developing improved male contraceptives.

It should be noted that collecting a good number of testicular samples for research is difficult and the high cost of mass-scale transcriptomic techniques becomes another obstacle. In fact, even in the case of microarray studies on human testes, less number of samples have been used. There could be socioeconomic reasons behind these discrepancies. The number of samples available for research on NOA seems to be less in many cases, including countries where population growth has been very high and research towards contraception is expected to be a priority. The prevalence of male infertility is also probably the same in such countries compared to the global trend. While the range of testis-donors used for NOA has been usually less than 10 in reports from China (e.g., 36), researchers from some of the developed countries seem to have managed collecting higher number (16 to 47) of samples (e.g., 37, 11). Transcriptome level studies on testis of Indian NOA patients have not been reported. Only recently a mass scale methodology study has been reported using the testis tissue of NOA patient-donors from India (38) with 30 samples. Hopefully, increased interests and collaborations can prompt physicians and researchers to take up studies with a higher number of samples in such developing countries as well, in the near future.

In any case, all applied research towards diagnostics and treatment of male infertility, as well as male contraceptive development, requires a clear understanding of the spermatogenesis itself. The current results can help to identify several new molecules associated with the NOA and/or normal spermatogenesis. Interestingly, some of the very well-studied genes, as indicated by current literature analysis, were not known to be associated with NOA so far. Some of the reasons for this may be the limitations in earlier microarray based transcriptomic studies on the NOA transcriptomics in terms of (a) the probe selections; (b) lesser number of genes covered in the earlier microarray-based mass scale studies; and (c) consideration of alternatively spliced forms in the current study. RNA-seq approach, on the contrary, is known to cover all genes and identify specific transcripts associated with conditions of interest.

The RT-qPCR experiments in the current study served two purposes: testing the biomarker candidature status of each transcript and validation of the RNA-sequencing experiments. A lower portion of top-scoring transcripts was expected to show differential expression in every NOA sample for two reasons: a) Given the complexity of primer regions across transcripts that would ensure unique regions and the varying coverage of exons by reads, certain primers may not work as per expectation with all samples. In fact, 12 primer-pairs corresponding to 12 transcripts never worked. b) Compared to only 8 samples used for NGS experiments, better coverage of diversity of NOA subtypes is likely to exist among all the (eighteen) NOA samples used across NGS and RT-qPCR experiments. Nevertheless, the overall trend in expression level seen in NGS experiments was not contradicted by the trend observed via RT-qPCR experiments - particularly for down-regulated transcripts. Thus, the latter experiments validated the reliability of RNA-seq data. It should be noted that the results of functional analysis and comparison with known markers from the literature also strongly support the reliability of the RNA-seq data.

Serious efforts to biocurate and compile genes associated with human spermatogenesis and NOA have not been reported earlier. The current study includes reasonable biocuration and data mining efforts. Though literature search, AnnoKey and GenCLIP, and GO term analysis have been used here for compiling, a more complete list of known genes, a more thorough biocuration via literature search, particularly involving multiple search engines (39), is still warranted to create a comprehensive list of key genes from literature. It should be noted that while our approach has focused more on ensuring that the ‘novelty’ of association by RNA-seq analysis is being reported correctly, there is a chance that the GenCLIP and AnnoKey results resulted in a few false positives as the mere co-occurrence of keywords has been considered positive.

Current data mining revealed that some of the genes found associated with NOA via RNA-seq experiment and analysis were also cited more frequently earlier in the context of spermatogenesis and NOA condition as indicated above. About 47 and 234 genes were already found to be associated with NOA and spermatogenesis, respectively, among the top 500 genes. While ‘the strength of association with NOA’ and the corresponding ‘specific transcript isoform’ are being identified for the first time for 4 of the 16 genes, the NOA association of the remaining 12 genes are being reported at the gene level also for the first time. Further, while two genes were cited in 80 and 29 research papers in the context of spermatogenesis, some others were cited up to 7 times and/or had spermatogenesis related GO terms assigned to them. Interestingly, none of the data mining approaches indicated any known association with spermatogenesis or NOA for 5 of the 16 short-listed candidate markers. Thus, probably the use of the StA-score-based ranking can help to identify novel genes mainly based on the strength of association while avoiding an over-emphasis on known genes, which is a common criteria to short-list genes after the first level of identification of disease-associated genes, which in turn is based on fold changes following a basic P-value cut off. This is similar to the approach we followed earlier for meta-analysis of microarray data, where consistency in expression patterns across the studies is given more weight than the fold change and P-values (40). The three trans-splicing events found to be promising candidate markers for NOA, seem to be novel for any condition.

Though the StA scoring seems useful in general, it should be noted that in the current study, the ranking based on StA was almost similar to the ranking based on only the fold-change for the top- 250 down-regulated genes. The rankings by these two methods produced different results for up-regulated transcripts and genes, and the bigger lists for down-regulated genes (e.g., top 3000 transcripts). Though useful in the current case, the specific advantages of this novel method should be further tested in more cases.

Irrespective of the sub-type of NOA, most patients are likely to have no fully functional sperm in the testis. Hence, it was expected that several specific functions related to spermatozoa, such as sperm capacitation, acrosome reaction and sperm motility would be suppressed. But, we did not expect suppression or impairment of spermatid development over-represented among genes down-regulated in ‘all’ NOA patients. This is because a significant number of NOA patients could have spermatids and/or sperm (e.g., 41, 42). It is less likely that all sequenced patients would have a pre-meiotic arrest in spermatogenesis. There is a possibility that genes picked in the GO enrichment analysis, are the ones commonly down-regulated in patients with ‘imperfectly developed spermatids’ as well as those with no spermatids. If this is true, even when spermatids exist in NOA patients they may not be fully functional in such NOA patients. This is also supported by a lack of proper spermiogenesis in most NOA patients and/or frequent failure of spermatid/sperm injections during IVF in NOA cases.

Cell-type-specific enriched expression of genes is critical and this has been addressed well (20) and can help to explore cell-type specificity of key genes. Of the top 250 genes down-regulated with high StA score in the current study, 148 genes were common to the significant cell-type-enriched list from Wang et al, and 133 of these were enriched in spermatids. On the contrary, only 11 of the top 250 up-regulated genes common to the significant cell-type-enriched list reported by single-cell sequencing earlier, and 10 of these 11 were enriched in spermatids.

Apart from obvious GO terms for broad biological processes, such as spermatogenesis, spermatid development, etc, many other specific terms were also found to be enriched among the 3,000 down-regulated genes in the current study, which have also been reported earlier to be key processes associated with NOA (e.g., 11, 43, 44). Though there were many common observations among the functions and pathways over-represented among genes up- or down-regulated as per earlier studies, a few new observations were also noted. Even though piRNA metabolic process was reported to be enriched among NOA-down-regulated genes in an integrated analysis of miRNAs and mRNAs (45), other transcriptomic analysis did not report piRNA metabolic process and ubiquitination, which were enriched among down-regulated genes in the current study. But, germline mutations in the human Piwi gene in patients with azoospermia have been reported to prevent the ubiquitination and degradation of the corresponding protein (46). Global ubiquitination of H2A and H2B mediated by the Piwi-dependent RNF8 protein seems to be a key step in histone to protamine transition (47). Even though integrated data analysis has been made earlier (44), with increasing data availability, there is a scope to perform an in depth functional analysis of human and mammalian genes in the context of their potential roles in spermatogenesis and NOA. An optimal use of the testis-specific gene expression database (40) may also be useful in this regard.

Viral DNA is frequently detected in the reproductive system of normal as well as infertile donors (48, 49). Herpes virus has been detected in the semen of donors - including those with normal sperm (50, 51). Some studies have also indicated active viral gene expression. For example, Epstein-Barr virus infection pathway has been earlier shown to be represented by genes up-regulated in meiotic arrest and Sertoli-cell only sub-types of NOA (42). Active transcription of viral genomes, as suggested by current observations has also been indicated earlier by mass-scale studies (42) as well as a specific study on Thymidine Kinase from human herpesvirus 1 in infertile men (52). The current observations did not indicate involvement of a specific viral pathway, but suggested a general viral gene expression; viral transcription and translation processes were common functions among the up-regulated transcripts. We also made a few new observations: while several viral molecular functions / biological processes seem to be more common or promoted in the testes from NOA patients than those from control donors, an anti-viral mechanism showed a reverse trend. Considering the above-listed findings together, we raise new questions: While a few types of viruses may have been successful in invading testes, is there a failure in the testes of some of the NOA-patients to curb viral growth? Could this be one of the causes of NOA?

## Conclusions

In the current study, transcriptomic analysis of the testes from donors with Non-Obstructive Azoospermia (NOA) has been used to identify the genes and the corresponding alternatively spliced mRNA-isoforms associated with the disorder. These molecules have been hierarchically listed based on their strength of association with the NOA condition, which would also suggest the association with spermatogenesis in most cases, particularly those down-regulated in NOA. Such an association, of several genes and most of the alternatively spliced forms of mRNAs, is being reported for the first time. Based on the 100% consistency of differential expression observed across 18 NOA and 22 control samples, via RNA-seq and/or the RT-qPCR results, 16 mRNAs are being suggested as promising candidates for reliable diagnosis of NOA. Similarly, 3 chimeric transcripts have also been short-listed as potential diagnostic markers. If these 19 markers could be used in RT-qPCR assays, individually or in combination with each other, they could avoid the need for open surgeries for the detection of NOA. The functional analysis of the top few hundreds of differentially transcribed genes suggested possible suppression of some of the molecular functions such as ubiquitination and anti-viral mechanisms, while viral gene expression seems to be promoted, in the testes of NOA patients. The current observations hopefully prompt further studies to explore the potential role of testicular viral infections in the context of NOA. It may be a useful strategy to prioritize the top-ranking transcripts from the current study, or the corresponding genes and proteins while exploring biomarkers for the NOA subtypes. The specific role of top-ranking genes in spermatogenesis, for which NOA association has been identified for the first time, also needs to be explored.

## Acknowledgements

Research projects at IBAB are supported by the Department of IT, BT and S&T, Government of Karnataka, India. Staff members at the Bio-IT center at IBAB carried out the RNA-sequencing work.

## Author contributions

RNA extraction and quality assessments, primer designing, RT-PCR and RT-qPCR: BG & BK; RT-PCRs: MM; preliminary RNA-seq data analysis, other data mining and analysis: AKB, SD, BG & BK; biocuration: BG, KS, BK, OS & CN; sample clustering: CDS & BG; scripting for data analysis: BV; sample identification & biopsy collection: VSS; original study design, experimental plans, task assignments, and guidance to all team members in conducting experiments and analysis: AKK.

## Competing interests

AKK was also the founder director of a private limited company, Shodhaka Life Sciences Pvt. Ltd., during the research period, and AKB & SD have also been employees of this company. Dr. Vasan is affiliated to commercial health care organizations as well, as listed above. The intellectual property of the diagnostic application of short-listed potential molecules and primer-pairs, including the new control pairs, is being protected via a patent application by IBAB and the Ankur Andrology & Men’s Health.

